# Developing an empirical model for spillover and emergence: Orsay virus host range in *Caenorhabditis*

**DOI:** 10.1101/2021.12.10.472097

**Authors:** Clara L. Shaw, David A. Kennedy

**Author notes:** Corresponding Author: David A. Kennedy.

## Abstract

A lack of tractable experimental systems in which to test hypotheses about the ecological and evolutionary drivers of disease spillover and emergence has limited our understanding of these processes. Here we introduce a promising system: *Caenorhabditis* hosts and Orsay virus, a positive-sense single-stranded RNA virus that naturally infects *C. elegans*. We assayed the susceptibility of species across the *Caenorhabditis* tree and found 21 of 84 wild strains belonging to 14 of 44 species to be susceptible to Orsay virus. Confirming patterns documented in other systems, we detected effects of host phylogeny on susceptibility. We then tested whether susceptible strains were capable of transmitting Orsay virus by transplanting exposed hosts and determining whether they transmitted infection to conspecifics during serial passage. We found no evidence of transmission in 10 strains (virus undetectable after passaging), evidence of low-level transmission in 5 strains (virus lost between passage 1 and 5), and evidence of sustained transmission in 6 strains (including all 3 experimental *C. elegans* strains). Transmission was associated with host phylogeny and with viral amplification in exposed populations. Variation in Orsay virus susceptibility and transmission among *Caenorhabditis* species suggests that the system could be powerful for studying spillover and emergence.

## INTRODUCTION

Disease spillover and emergence can have catastrophic consequences for the health of humans and other species. For example, SARS-CoV-2 spilled over into human populations [1] and became pandemic, killing more than 5 million people when this study was published [2]. Moreover, the frequency of spillover events and the rate of new disease emergence has been increasing in the recent past [3], endowing urgency to the task of understanding drivers of spillover and the progression of emergence. Studies in wild systems with ongoing spillover have provided substantial insights into the spillover and emergence process [4–6], but experimental manipulation to test hypotheses in these systems can be impractical due to ethical and logistical concerns. Moreover, disease emergence is so rare that it typically can only be studied retrospectively. Therefore, it remains a challenge to understand what factors facilitate emergence and how evolution proceeds in emerging pathogens.

Spillover requires that pathogens have the opportunity and the ability to exploit a new host; emergence requires that this opportunity and ability persist through time [5,7]. Opportunity could arise if hosts share habitats or resources. Ability may arise through mutations or pre-exist due to pathogen plasticity or host similarity. Studies of natural spillover and emergence events have identified characteristics of pathogens, hosts, and their interactions that generally support the above. For example, pathogens that successfully spill over are likely to be RNA viruses with large host ranges [8,9]. Likewise, hosts with close phylogenetic relationships are more likely to share pathogens than more distantly related hosts [9–14]. In addition, geographic overlap between hosts is associated with sharing pathogens [12], meaning that changes in host population distributions that bring new species into contact could potentially promote spillover and emergence events [9,15–17].

Ecological factors (e.g. host densities, distributions, diversity, condition, and behavior) can promote or hinder spillover by modulating host exposure risk or host susceptibility [5,7]. Likewise, it is believed that ecological factors can promote or hinder emergence through the modulation of onward transmission in spillover hosts, which determines whether pathogens meet dead ends in novel hosts, transmit in stuttering chains, or adapt and persist [18–20]. Conclusively demonstrating the influence of ecological factors, however, requires experimental manipulation, and it has so far been difficult to perform such studies.

Experimental model systems have been essential for testing hypotheses about infectious disease biology [21–23]. Indeed, major discoveries in immunity, pathogenesis, and pathogen ecology and evolution come from model systems such as *Mus musculus* [24], *Drosophila melanogaster* [25], *Daphnia* species [21], *Arabadopsis thaliana* [26], and *Caenorhabditis elegans* [27]. These systems have important traits that make them amenable to experimentation: they are inexpensive, have fast generation times, and have simplified genetics since they are usually hermaphroditic, asexual, or inbred. In addition, experimental tools and knowledge have accumulated in these systems, lowering the barriers to novel findings. However, few model systems exist to study the ecology and evolution of disease spillover and emergence, and the systems that do exist lack key features known to drive disease dynamics (e.g. host behavior or transmission ecology). A perfect model system would have large host population sizes, naturally transmitting, fast-evolving pathogens (e.g. viruses), and multiple potential host species with variable susceptibility and transmission.

*Caenorhabditis* nematode species are appealing model host candidates. Indeed, *C. elegans* and various bacterial and microsporidian parasites are staples of evolutionary disease ecology [22,28]. Specifically, the trivial manipulation and sampling of laboratory host populations means that population-level processes like disease transmission and evolution can be observed, and the tractable replication of large populations makes possible the observation of rare events like spillover and emergence. However, until recently, there were no known viruses of any nematodes including *C. elegans*. That changed with the recent discovery of Orsay virus [29].

Orsay virus, a natural gut pathogen of *C. elegans*, is a bipartite, positive-sense, single-stranded RNA (+ssRNA) virus that transmits readily in laboratory *C. elegans* populations through the fecal-oral route [29]. This virus is an appealing model pathogen candidate since +ssRNA viruses have high mutation rates [30] and typically evolve quickly [31]. Moreover, since Orsay virus transmits between hosts in the lab, this system allows transmission itself to evolve, a critical component of emergence [19] that cannot be readily studied in other animal laboratory systems of disease emergence. To develop *Caenorhabditis* hosts and Orsay virus as a system for studying spillover and emergence, it is necessary to know the extent to which the virus can infect and transmit in non-*elegans Caenorhabditis* species. So far, such exploration been limited to one other species, *C. briggsae*, which was determined to be refractory to infection [29]. Notably, an ancestral virus likely crossed at least one host species boundary in the past since *C. briggsae* has been found to be susceptible to three related viruses [29,32,33].

To explore the suitability of the *Caenorhabditis*-Orsay virus system for studies of disease spillover and emergence, we first test a suite of *Caenorhabditis* species for susceptibility to Orsay virus, and then we test the extent to which susceptible host species can transmit the virus. For both traits (susceptibility and transmission ability), we test for effects of host phylogeny. Though host ranges of pathogens have been studied by infection assays (e.g. [34–37]) or by sampling infected hosts from natural systems (e.g. [11,38]), these studies do not typically distinguish between dead-end infections, stuttering chains of transmission, and sustained transmission. Therefore, to our knowledge, our study is the first to empirically link phylogeny with disease transmission dynamics in novel species following spillover.

## METHODS

### Susceptibility Assays

We assayed susceptibility of *Caenorhabditis* species to Orsay virus by measuring virus RNA in previously virus-exposed host populations using quantitative PCR (qPCR). We obtained 84 wild isolate strains belonging to 44 *Caenorhabditis* species (1-3 strains per species) from the *Caenorhabditis* Genetics Center (CGC) and from Marie-Anne Félix. We tested each strain for Orsay virus susceptibility using 8 experimental blocks (Table 1, Table S1). Species identities were confirmed by sequencing the small ribosomal subunit internal transcribed spacer ITS2 and/or by mating tests. For each *Caenorhabditis* strain, we initiated three replicate populations with five adult animals. For sexual species, we used five mated females, and for hermaphroditic species, we used five hermaphrodites. All populations were maintained on nematode growth medium (NGM) in 60 mm diameter plates with a lawn of bacterial food (lawns were seeded with 200 µL *E. coli* strain OP50 in Luria-Bertani (LB) broth and allowed to grow at room temperature for approximately 24 hours [39]). We exposed populations to virus by pipetting 3 µL of Orsay virus filtrate, prepared as described in [29], onto the center of the bacterial lawn. We determined the concentration of the filtrate to be 428.1 (95% CI: 173.4-972.3) x the median tissue culture infectious dose (TCID50) per µL (Supplement A) [40]. We maintained populations at 20°C until freshly starved (i.e. plates no longer had visible bacterial lawns). Depending on the strain, this took anywhere from 3 to 28 days (Table S1). While this meant that strains may have experienced variable numbers of generations, this method ensured that all the exposure virus was consumed. We collected nematodes from freshly starved plates by washing plates with 1,800 µL water and transferring suspended animals to 1.7 mL microcentrifuge tubes. We centrifuged tubes at 1000 x g for 1 minute to pellet nematodes. We removed the supernatant down to 100 µL (including the pellet of nematodes) and ‘washed’ external virus from nematodes by adding 900 µL of water and removing it 5 times, centrifuging at 1000 x g for 1 minute between each wash. After the five washes, we lysed the nematodes by transferring the nematode pellet along with 500 µL water to 2 mL round-bottom snap cap tubes, adding approximately 100 µL of 0.5 mm silica beads, and shaking in a TissueLyser II (Qiagen) for 2 minutes at a frequency of 30 shakes per second. We then removed debris with two centrifugation steps of 17,000 x g for 5 minutes, each time keeping the supernatant and discarding the pellet. Samples were stored at −80 °C.

**Table 1.** Strains assayed for susceptibility to Orsay virus with the number of replicates processed in each block. When strains were assayed in multiple blocks, replicate numbers are given in the respective order of the blocks. Strains were acquired from the *Caenorhabditis* Genetics Center (University of Minnesota) and from Marie-Anne Felix (IBENS).

| Strain | Species | Block | Number of Replicates | Strain | Species | Block | Number of Replicates |
| --- | --- | --- | --- | --- | --- | --- | --- |
| JU1199 | *C. afra* | 2 | 3 | JU2613 | *C. portoensis* | 7 | 3 |
| JU1198 | *C. afra* | 4 | 3 | JU2745 | *C. quiockensis* | 2 | 3 |
| JU1593 | *C. afra* | 7 | 3 | MY28 | *C. remanei* | 2 | 3 |
| NIC1040 | *C. astrocarya* | 3 | 1 | PB206 | *C. remanei* | 6 | 3 |
| QG704 | *C. becei* | 2 | 3 | JU1082 | *C. remanei* | 6 | 3 |
| SB280 | *C. brenneri* | 1 | 3 | JU1201 | *C. sinica* | 1 | 3 |
| SB129 | *C. brenneri* | 6 | 3 | JU4053 | *C. sinica* | 4 | 3 |
| LKC28 | *C. brenneri* | 6 | 3 | JU1202 | *C. sinica* | 6 | 3 |
| JU1038 | *C. briggsae* | 1,2,3<sup>1</sup> | 3,3,3 | JU2203 | *C. sp. 8* | 5 | 2 |
| EG4181 | *C. briggsae* | 6 | 3 | QG555 | *C. sp. 24* | 3 | 3 |
| ED3083 | *C. briggsae* | 6 | 3 | JU2867 | *C. sp. 24* | 5,7 | 1,3 |
| JU1426 | *C. castelli* | 3,7 | 3,3 | JU2837 | *C. sp. 24* | 6 | 3 |
| JU1333 | *C. doughertyi* | 1 | 3 | ZF1092 | *C. sp. 25* | 3 | 3 |
| JU1328 | *C. doughertyi* | 4 | 3 | QX2263 | *C. sp. 27* | 1,3 | 2,3 |
| JU1331 | *C. doughertyi* | 5 | 3 | DF5152 | *C. sp. 30* | 3 | 3 |
| DF5112 | *C. drosophilae* | 3 | 3 | NIC1070 | *C. sp. 43* | 2 | 3 |
| GXW1 | *C. elegans* | 6 | 3 | JU4050 | *C. sp. 62* | 5 | 3 |
| JU1401 | *C. elegans* | 6 | 3 | JU4045 | *C. sp. 62* | 7 | 3 |
| ED3042 | *C. elegans* | 6 | 3 | JU4056 | *C. sp. 63* | 6 | 3 |
| NIC113 | *C. guadaloupensis* | 1 | 3 | JU4061 | *C. sp. 64* | 6 | 3 |
| EG5716 | *C. imperialis* | 3 | 3 | JU4087 | *C. sp. 65* | 4 | 3 |
| JU1905 | *C. imperialis* | 7 | 3 | JU4093 | *C. sp. 65* | 5 | 3 |
| NKZ35<sup>2</sup> | *C. inopinata* | 3 | 3 | JU4092 | *C. sp. 65* | 5 | 3 |
| QG122 | *C. kamaaina* | 2 | 3 | JU4094 | *C. sp. 66* | 4 | 3 |
| VX80 | *C. latens* | 1 | 3 | JU4096 | *C. sp. 66* | 4 | 3 |
| JU3325 | *C. latens* | 4 | 3 | JU4088 | *C. sp. 66* | 4 | 3 |
| JU724 | *C. latens* | 5,7 | 1,3 | SB454 | *C. sulstoni* | 2 | 3 |
| JU1857 | *C. macrosperma* | 2 | 3 | JU2774 | *C. tribulationis* | 1 | 3 |
| JU1865 | *C. macrosperma* | 5 | 3 | JU2776 | *C. tribulationis* | 5 | 3 |
| JU1853 | *C. macrosperma* | 7 | 3 | JU2775 | *C. tribulationis* | 5 | 3 |
| JU2884<sup>3</sup> | *C. monodelphis* | 8 | 3 | JU1373 | *C. tropicalis* | 1 | 3 |
| JU1667<sup>3</sup> | *C. monodelphis* | 8 | 3 | JU1428 | *C. tropicalis* | 2 | 3 |
| JU1325 | *C. nigoni* | 1,2,3 | 2, 1, 3 | JU2469 | *C. uteleia* | 2 | 3 |
| JU2617 | *C. nigoni* | 4 | 3 | JU2458 | *C. uteleia* | 4 | 3 |
| EG5268 | *C. nigoni* | 6 | 3 | JU1968 | *C. virilis* | 3 | 3 |
| JU1825 | *C. nouraguensis* | 1 | 3 | JU2758 | *C. virilis* | 5 | 3 |
| JU1833 | *C. nouraguensis* | 5 | 3 | NIC564 | *C. waitukubuli* | 1 | 3 |
| JU1854 | *C. nouraguensis* | 6 | 3 | JU1873 | *C. wallacei* | 1 | 3 |
| QG702 | *C. panamensis* | 2 | 3 | EG6142 | *C. yunquensis* | 3 | 3 |
| JU2770 | *C. parvicauda* | 7 | 3 | JU2156 | *C. zanzibari* | 1 | 3 |
| EG4788 | *C. portoensis* | 1 | 3 | JU3236 | *C. zanzibari* | 6 | 3 |
| JU3126 | *C. portoensis* | 5 | 3 | JU2161 | *C. zanzibari* | 7 | 3 |

<sup>1</sup>JU1038 was included in the first three blocks as a type of negative control since a previous study found that *C. briggsae* was not susceptible. We discontinued this practice given the number of strains we needed to test.
<sup>2</sup>Strain NKZ35 was maintained at 23°C according to *Caenorhabditis* Genetics Center recommendation.
<sup>3</sup>Populations were initiated with 12 juvenile animals due to challenges rearing animals with standard methods.

We used qPCR to measure viral RNA in these samples. Primers and probe were: Forward: GTG GCT GTG CAT GAG TGA ATT T, Reverse: CGA TTT GCA GTG GCT TGC T, Probe: 6-FAM-ACT TGC TCA GTG GTC C-MGB. We performed 10 µL reactions composed of 1.12X qScript XLT One-Step RT-qPCR ToughMix (Quantabio), 200 nM each of forward and reverse primers and probe, and 2 µL of sample. Reaction conditions were: 50 °C (10 min), 95 °C (1 min), followed by 40 cycles of 95 °C (3 sec), 60 °C (30 sec). Assays were run on a 7500 Fast Real-Time qPCR System (Thermo Fisher Scientific, Applied Biosystems). Cycle threshold (Ct) values were determined using the auto-baseline and auto-threshold functions of the 7500 Fast Real-Time software (Thermo Fisher Scientific, Applied Biosystems).

Each experimental block also contained five sets of controls and benchmarks (Table 2): a negative control where virus was never added (control 1), two positive controls where strains with known susceptibilities were exposed (control 2, strain N2: mean(Ct)=15.7, sd(Ct)=2.0; control 3, strain JU1580: mean(Ct)=12.7, sd(Ct)=2.2), a benchmark to determine a Ct threshold for infection (benchmark 4: mean(Ct)=38.4, sd(Ct)=2.6), and a benchmark that gives a conservative Ct threshold for viral replication (benchmark 5: mean(Ct)=22.0, sd(Ct)=0.6). Species were considered susceptible if at least one replicate population amplified virus to levels higher than our infection threshold (one standard deviation more virus than the maximum value of benchmark 4 across all blocks which translates to Ct<29.5).

**Table 2.** Description of controls and benchmarks included in triplicate in each of the 8 blocks of the susceptibility assays.

| Control/benchmark | Description | Type |
| --- | --- | --- |
| 1 | Laboratory *C. elegans* strain N2 exposed to 3 μL water | Negative control |
| 2 | Laboratory *C. elegans* strain N2 exposed to 3 μL Orsay virus filtrate | Positive control |
| 3 | Highly susceptible *C. elegans* strain JU1580 exposed to 3 μL of Orsay virus filtrate | Positive control |
| 4 | 3 μL Orsay virus filtrate pipetted on the center of bacterial lawn with no nematodes | Threshold<sup>a</sup> |
| 5 | 3 μL Orsay virus filtrate added directly to 497 μL water, yielding the final extraction volume for experimental populations. | Threshold<sup>b</sup> |

<sup>a</sup>The purpose of this benchmark was to quantify exposure virus remaining in samples after 5 rounds of washing.
<sup>b</sup>The purpose of this benchmark was to quantify the maximum amount of virus that could be present without replication (i.e. total amount of virus added to each plate).

### Transmission Assays

We conducted transmission assays for all strains where at least one replicate population was determined to be infected in our susceptibility assay. First, three replicate populations were initiated as above and exposed to 3 µL of virus filtrate. At the same time, we initiated three replicate positive control populations of *C. elegans* laboratory strain N2 exposed to 3 µL virus filtrate and three replicate negative control populations of N2s exposed to 3 µL of water. When populations were recently starved, 20 adult nematodes (mated females for sexual species or hermaphrodites for hermaphroditic species) were chosen at random and passaged to virus-free plates with fresh food (*E. coli* strain OP50 lawns prepared as above). Remaining animals were washed from the starved plates, virus was extracted, and viral RNA quantified via qPCR as above (Table S2). We passaged each replicate line 5 times, or until there was no detectable viral RNA by qPCR. Controls were passaged 5 times regardless of virus detection.

We assigned each passage line a transmission score of 0, 1, 2, or 3 based on detection of viral RNA through the passages. A value of 0 was assigned when viral RNA was not detected in the exposure population; a value of 1 was assigned when viral RNA was detected in the exposure population but not in the first passage population; a value of 2 was assigned when viral RNA was detected in the first passage population but became undetectable on or before the fifth passage population; and a value of 3 was assigned when viral RNA was still detectable in the fifth passage population.

### Statistical Analysis

To test for phylogenetic effects, we fit Bayesian phylogenetic mixed effects models to the susceptibility and transmission data using the ‘MCMCglmm’ package [35,41,42] in R [43]. For these models, we used the most recent published phylogeny of *Caenorhabditis* [44]. We rooted the phylogeny with *Diploscapter pachys* as the outgroup and constrained it to be ultrameric (i.e. tips are all equidistant from the root) using the ‘chronopl’ function in the ‘ape’ package [45] with a smoothing parameter of 1. Since our susceptibility data are binomial, we fit them using logistic regression with a logit link. In practice this was achieved by setting family to ‘multinomial2’. Our transmission data are continuous, and we fit them using linear regression by setting family to ‘gaussian’. Data from controls and benchmarks were excluded from the analysis. For both the susceptibility and transmission data, all models included a random effect of species and all transmission models also included a random effect of strain. These random effects were included to prevent pseudo-replication. Other factors were included or excluded as described below. For the susceptibility data, our most complicated model included effects of phylogenetic distance to the native host *C. elegans* (calculated by the ‘cophenetic.phylo’ function in ‘ape’ [45]) and phylogenetic distance between pairwise sets of species (calculated by the ‘inverseA’ function in ‘MCMCglmm’ [41,46]). Note that ‘inverseA’ calculates the inverse relatedness matrix (i.e. the inverse of the matrix that contains the time from the root to the common ancestor of each species pair), but we refer to this metric as “phylogenetic distance between pairwise sets of species” for simplicity. For the transmission data, our most complicated model included these effects and an additional effect of viral amplification in the primary exposure population measured as Ct. Phylogenetic distance from *C. elegans* and viral amplification in the primary exposure population were treated as fixed effects, and phylogenetic distance between pairwise sets of species was treated as a random effect. We generated a suite of nested models that included all possible combinations of including or excluding these effects (Table 3, Table 4).

**Table 3.** Models compared for analysis of susceptibility patterns. All models included an intercept. The random effect of species is retained in all models to avoid pseudo-replication.

| Model | ΔDIC | DIC weight |
| --- | --- | --- |
| Suscep. ~ **fixed** = phylo. dist., **random** = pairwise phylo. dist. + species | 0 | 0.486 |
| Suscep. ~ **fixed** = phylo. dist., **random** = species | 1.121 | 0.277 |
| Suscep. ~ **fixed** = **random** = pairwise phylo. dist. + species | 2.189 | 0.163 |
| Suscep. ~ **fixed** = **random** = species | 3.761 | 0.074 |

‘phylo. dist’ indicates the effect of phylogenetic distance from *C. elegans* whereas ‘pairwise phylo. dist.’ indicates the effect of phylogenetic distance between species pairs.

**Table 4.** Models compared for analysis of transmission scores. All models included an intercept. Random effects of species and strain are retained in all models to avoid pseudo-replication.

| Model | ΔDIC | DIC weight |
| --- | --- | --- |
| Trans. ~ **fixed** = Ct + phylo. dist., **random** = pairwise phylo. dist. + species + strain | 0 | 0.269 |
| Trans. ~ **fixed** = Ct , **random** = pairwise phylo. dist. + species + strain | 0.533 | 0.206 |
| Trans. ~ **fixed** = Ct + phylo. dist., **random** = species + strain | 0.585 | 0.201 |
| Trans. ~ **fixed** = Ct , **random** = species + strain | 0.790 | 0.181 |
| Trans. ~ **fixed** = phylo. dist., **random** = pairwise phylo. dist. + species + strain | 3.942 | 0.038 |
| Trans. ~ **fixed** = **random** = species + strain | 4.086 | 0.035 |
| Trans. ~ **fixed** = **random** = pairwise phylo. dist. + species + strain | 4.091 | 0.035 |
| Trans. ~ **fixed** = phylo. dist., **random** = species + strain | 4.112 | 0.034 |

‘Ct’ indicates viral amplification on primary exposure plates. ‘phylo.dist’ indicates the effect of phylogenetic distance from *C. elegans* whereas ‘pairwise phylo. dist.’ indicates the effect of phylogenetic distance between species pairs.

We used the MCMCglmm default priors for fixed effects (normal distribution with mean = 0 and variance = 10^8^) and parameter expanded priors for random effects that result in scaled multivariate F distributions with V=1, nu=1, alpha.mu=0, alpha.V=1000 [47]. Residuals were assigned inverse Wishart priors with V=1 n=0.002 [41]. We ran models for 100,000,000 iterations with a burn in of 300,000 and thinning interval of 10,000. We visualized traces to affirm convergence of MCMC chains.

We used the deviance information criterion (DIC) to describe the relative support of models and to understand the importance of parameters [48]. We calculated DIC weights for each model, each parameter, and the phylogenetic parameters combined [49]. The DIC weight of a model, calculated as 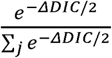 where *j* is the set of all models, gives the relative support for each model. Similarly, the DIC weight of a parameter, calculated as 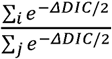 where refers to the set of models that includes a given parameter and *j* is the set of all models, is the posterior probability that a given factor is included in the ‘true’ model assuming the ‘true’ model has been designated. Thus, parameters with DIC weights greater than 0.5 are more likely than not to be included in the ‘true’ model.

## RESULTS

### Susceptibility Assays

In our assays of host susceptibility to Orsay virus, we identified 21 susceptible *Caenorhabditis* strains of the 84 experimental strains tested. These included three (non-control) strains of *C. elegans* (note that one of these strains JU1401 had been previously documented to be susceptible [50]) and 18 strains belonging to 13 other species. The strains were distributed broadly across the *Caenorhabditis* phylogenetic tree and in species that do not currently have a well determined phylogenetic placement (Figure 1). In total, we found that Orsay virus is capable of infecting hosts from at least 14 of 44 *Caenorhabditis* species.

**Figure 1.**
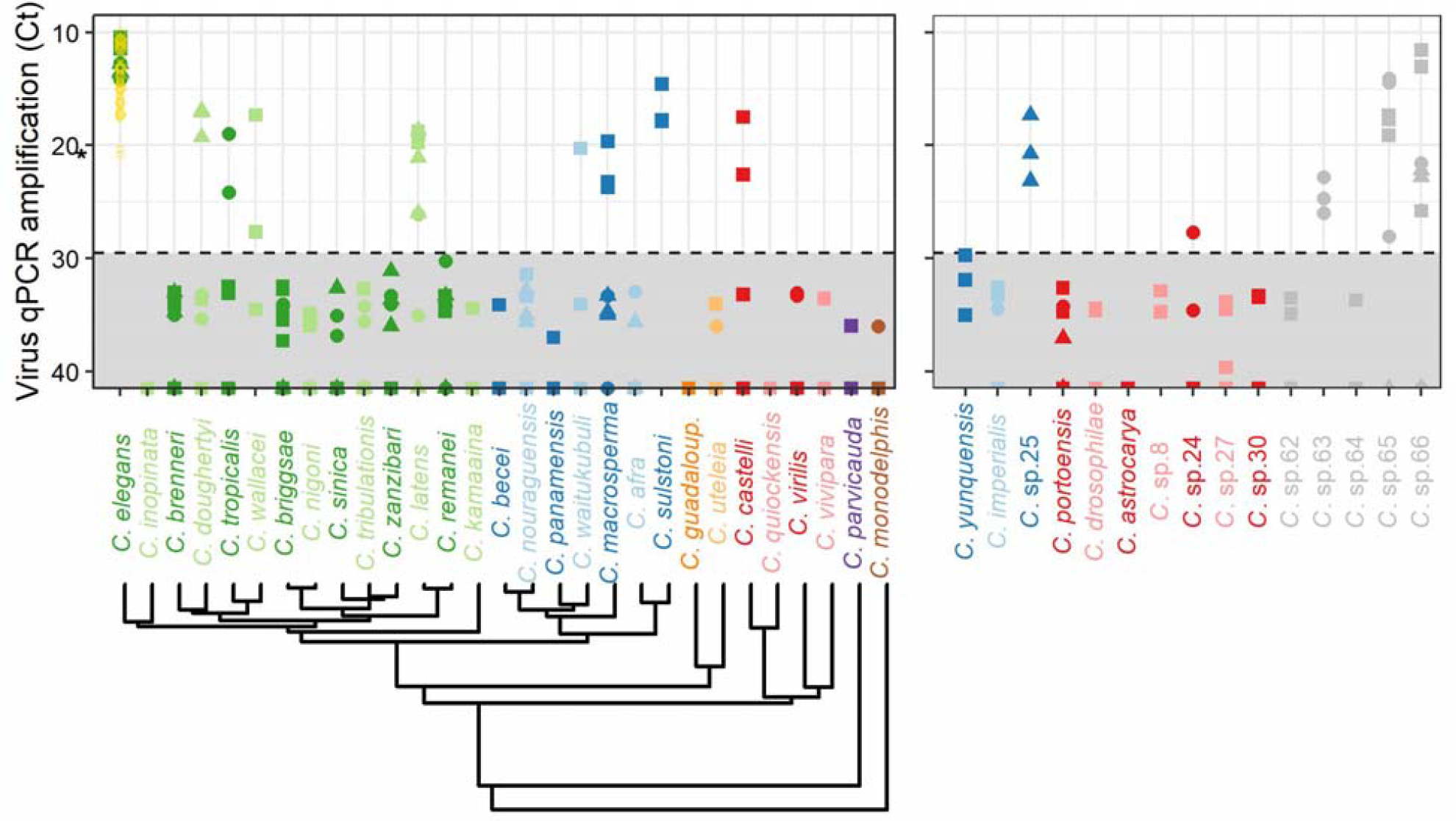
Species across the *Caenorhabditis* phylogeny are susceptible to Orsay virus (i.e. Ct values smaller than the infection determination cut off (dashed line, see methods). Note that smaller Ct values imply more virus). The asterisk on the left side of the y-axis shows the Ct value from ‘benchmark 5’ for the sample with the most detectable virus (Table 2). The phylogeny (bottom left) is pruned from [44]. Many species currently have uncertain phylogenetic placement (right). Species for which a clade is hypothesized are color-coded accordingly. These hypotheses were obtained from [53]. However, clades are unknown for *C*. sp. 62, *C*. sp. 63, *C*. sp. 64, *C*. sp. 65, *C*. sp. 66. Shapes indicate different strains within a species, colors differentiate clades, but are otherwise only varied to aid visualization. Open gold circles and diamonds indicate Ct values for positive controls (‘control 2’ and ‘control 3’ plates respectively; Table 2).

Our statistical analysis uncovered the importance of host phylogeny in explaining differences in susceptibility. Our best model included both phylogenetic effects tested: phylogenetic distance from *C. elegans* and phylogenetic distance between pairwise sets of species (Table 3). The model lacking these phylogenetic effects had a ΔDIC of 3.761 demonstrating support for the importance of phylogenetic effects [51,52]. We also computed DIC weights of parameters to show the relative importance of each on model fit. Distance from *C. elegans* had a weight of 0.763 and pairwise phylogenetic distance between sets of species had a weight of 0.648. Since both weights are greater than 0.5, each phylogenetic effect is more likely than not to be included in the ‘true’ model. Moreover, models that included at least one of these phylogenetic effects had a weight of 0.926, demonstrating very strong support for phylogenetic effects on susceptibility.

### Transmission Assays

The primary exposure populations (passage 0) in our transmission assay were treated nearly identically to populations in our susceptibility assay. As an internal control, we thus note high concordance between Ct measures in both assays (correlation coefficient = 0.85). Most replicates of *C. elegans* strains as well as positive control replicates (*C. elegans* strain N2) maintained high levels of virus through five passages (Figure 2). However, virus was lost in 1 out of 3 control replicates in both blocks; in retrospect, this is unremarkable since the N2 strain used for controls is known to be more resistant to Orsay virus than many other *C. elegans* strains [29]. Non-*elegans* strains did not transmit the virus as well in most cases. Virus was undetectable in the first passage population in all replicates of *C. doughertyi, C. wallacei, C. latens* strain JU3325, *C. waitukubuli, C*. sp. 25, *C. castelli, C*. sp. 24, *C*. sp. 63, and *C*. sp. 66 strains JU4088 and JU4096. Virus was also undetectable in the first passage population in one or two replicates of *C. tropicalis, C. latens* strain 724, *C. macrosperma, C. sulstoni, C*. sp. 65 strain JU4087, and *C*. sp. 66 strain JU4094. Virus was maintained for 1-4 passages in at least one replicate of strains of *C. tropicalis, C. latens* strain VX80, *C. macrosperma, C. sulstoni, C*. sp. 65 strains JU4093 and JU4087, and *C*. sp. 66 strain JU4094. Virus was detectable through the 5^th^ passage in four non-*elegans* replicates belonging to three strains of different species: 1 replicate of *C. sulstoni* strain SB454, 1 replicate of *C. latens* strain JU724, and 2 replicates of *C*. sp. 65 strain JU4093 (Figure 2).

**Figure 2.**
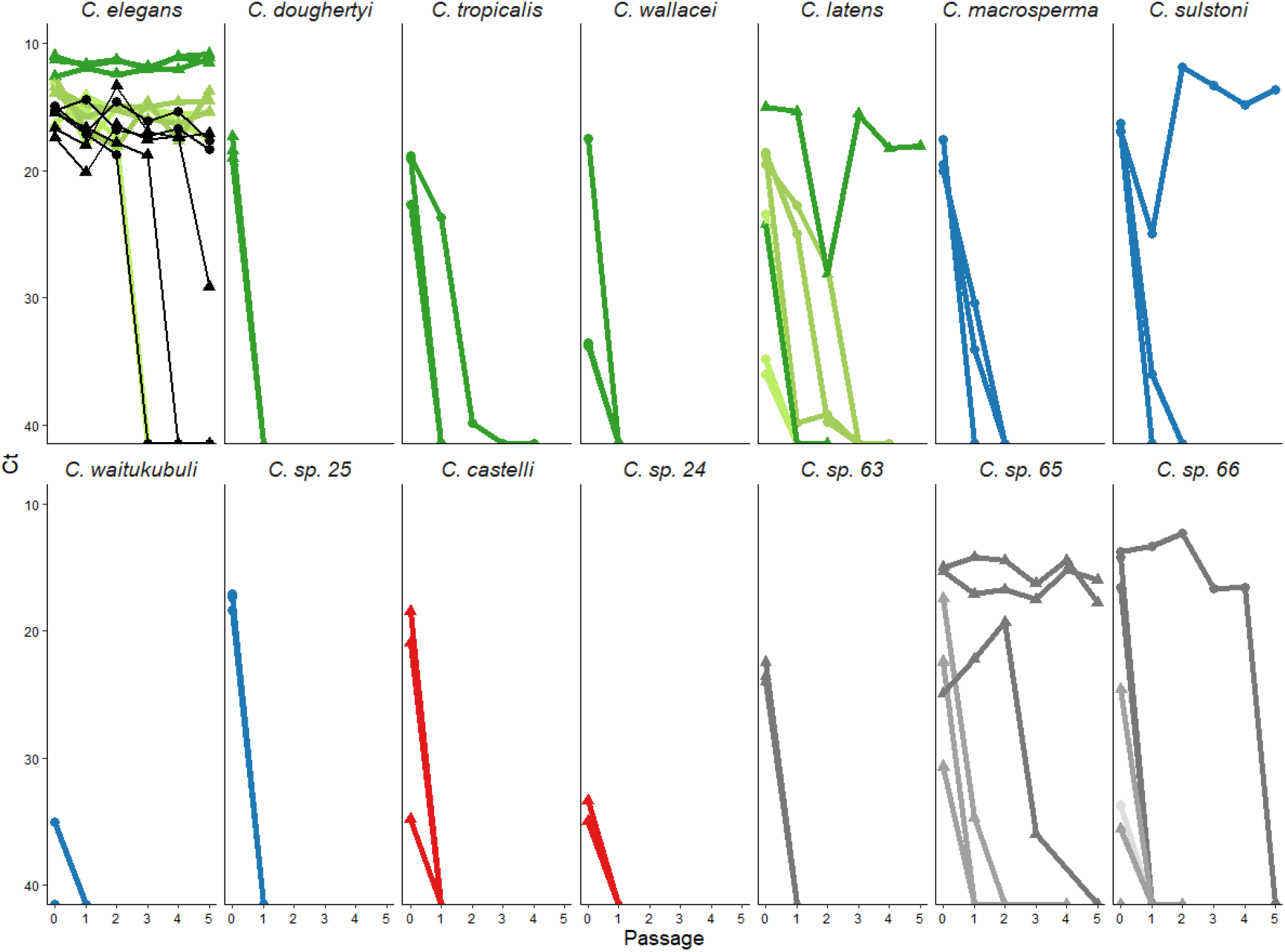
Orsay virus persisted to different extents when susceptible hosts were sequentially passaged to virus-free plates. “Passage 0” denotes the primary exposure population. This experiment was carried out in two blocks indicated by shape (circle=block 1, triangle=block 2). N2 controls were present in both blocks, shown in black. Colors match color-coded phylogeny in Figure 1. Shades represent different strains within a species: *C. elegans* GXW1 (dark green), ED3042 (medium green), JU1401 (light green); *C. doughertyi* JU1331; *C. tropicalis* JU1428; *C. wallacei* JU1873; *C. latens* JU724 (dark green; one of the three replicate lines was removed from analysis due to bacterial contamination), VX80 (medium green), JU3325 (light green); *C. macrosperma* JU1857; *C. sulstoni* SB454; *C. waitukubuli* NIC564; *C*. sp. 25 ZF1092, *C. castelli* JU1426; *C*. sp. 24 JU2837; *C*. sp. 63 JU4056; *C*. sp. 65 JU4093 (dark gray), JU4087 (medium gray); *C*. sp. 66 JU4094 (dark gray), JU4088 (medium gray), JU4096 (light gray).

As with the susceptibility data, we again identified factors associated with differences in transmission through model analysis. Our best model again included phylogenetic effects of distance from *C. elegans* and phylogenetic distance between pairwise sets of species. This model additionally included an effect of viral amplification (Ct) in primary exposure populations (Table 4), which was correlated with phylogenetic distance from *C. elegans* (correlation coefficient = 0.461). DIC weights were as follows: amplification (Ct) in primary exposure populations = 0.858, phylogenetic distance from *C. elegans* = 0.542, pairwise phylogenetic distance between sets of species = 0.548. Models including at least one of the phylogenetic effects had a weight of 0.784. These weights indicate strong support for an effect of viral amplification in primary exposure populations and at least some support for each phylogenetic effect in explaining transmission ability.

## DISCUSSION

In our study examining the host range of Orsay virus, we determined that at least 13 *Caenorhabditis* species in addition to *C. elegans* are susceptible and that hosts varied in their ability to transmit the virus. Specifically, we found 21 susceptible *Caenorhabditis* strains (including 3 out of 3 *C. elegans* strains) out of 84 tested strains belonging to 44 species. When susceptible strains were assayed for transmission ability, 10 strains were dead-end hosts in all replicates, and 6 strains (3 *C. elegans* strains, 1 *C. sulstoni* strain, 1 *C. latens* strain, and 1 *C*. sp. 65 strain) showed virus persistence for at least five passages in at least one replicate. The remaining 5 susceptible strains showed stuttering chains of transmission in at least one replicate. Both susceptibility and transmission ability were associated with two phylogenetic effects: distance from *C. elegans* and phylogenetic distance between pairwise sets of species. Transmission ability was also positively associated with viral amplification in primary exposure populations. Overall, we argue that this study primes the *Caenorhabditis*-Orsay virus system to be valuable for experimental studies on the ecology and evolution of pathogen spillover and emergence.

Replicating findings from several other experimental studies of host range [34–36], we found evidence of phylogenetic effects on susceptibility. Host species more closely related to the native host *C. elegans* were more likely to be susceptible to infection, and closely related hosts had more similar susceptibilities regardless of their relationship to the native host. These patterns may arise because closely related hosts likely have similar receptors, pathogen defenses, and within-host environments [10]. We expect that the importance of phylogenetic effects would only become more readily detectable if our unplaced *Caenorhabditis* species were placed on the phylogeny, since their lack of placement cost us statistical power. Importantly, we recovered an effect of phylogenetic distance from *C. elegans* even though few species are closely related to *C. elegans* (Figure 1). We hypothesize that the statistical support for this phylogenetic effect would become stronger if this work were repeated with related viruses of *C. briggsae*, which is a member of a clade with more closely related species.

We also found detectable effects of phylogeny on transmission ability. Although patterns consistent with a phylogenetic effect on transmission have been identified [10,35,54], to the best of our knowledge, this study is the first to empirically document such a pattern. In comparison to susceptibility, however, the association between phylogeny and transmission ability had weaker statistical support. This reduction in statistical support may have resulted from the small number of hosts tested, since we were only able to assay transmission in susceptible strains. Moreover, the susceptible species were less well distributed across the phylogenetic tree than random (i.e. the mean distance from *C. elegans* for strains in this assay was 0.149 and ranged from 0 to 0.419, while the mean distance from *C. elegans* across all strains in the susceptibility assay was 0.220 and ranged from 0 to 0.794). In addition, the moderate correlation between phylogenetic distance from *C. elegans* and our other focal fixed effect, viral amplification in primary exposure populations, may have made a phylogenetic distance effect more difficult to detect.

The strongest predictor of transmission ability in our study was viral amplification in primary exposure populations. We can imagine at least three reasons why amplification in primary exposure populations may matter for transmission. First, high levels of viral amplification may be indicative of some level of “pre-adaptation”, the ability to infect and transmit among novel hosts before additional evolutionary changes [55]. Indeed, the correlation between viral amplification in primary exposure populations with phylogenetic distance from *C. elegans* is consistent with this idea. Second, if hosts can shed the virus, high levels of viral amplification may expose conspecifics to higher doses, which could increase infection prevalence. If this was the case in our experiment, animals passaged from primary exposure populations with more viral amplification may have been more likely to have been infected. Third, larger virus populations may harbor more genetic variation, increasing opportunities for adaptive evolution that could maintain persistence of the virus in the spillover host. Indeed, evolutionary rescue theory has shown that larger populations are more likely to persist in comparison to smaller ones [56].

Here we have documented spillover and transmission of Orsay virus in *Caenorhabditis* hosts. It is important to note, however, that the patterns we see with our susceptibility and transmission assays may not fully predict spillover and emergence patterns among *Caenorhabditis* hosts in the wild.

Exposure risk is a key determinant of spillover and emergence [7], but in our experiments, we exposed all hosts equally. Orsay virus exposure risk for *Caenorhabditis* species in nature is unknown since we know little about the distributions of *Caenorhabditis* species and their viruses [57,58]. The two host species that have been most extensively studied in the wild, *C. elegans* and *C. briggsae*, do have overlapping distributions [59], but appear to be refractory to each other’s viruses [29]. However, the fact that three viruses related to Orsay virus have been found in *C. briggsae* [29,32,33] suggests that at least one host jump has occurred in the past, since the viruses appear to be much more closely related [33] than *C. briggsae* and *C. elegans* [60].

*C. elegans* has long been used as a model system to study infectious disease [22]. We argue that the *Caenorhabditis-*Orsay virus system will be useful for studying virus spillover and emergence since the system has many attractive features, including large populations, short experimental timelines, replicable experimental manipulations, natural transmission, and related hosts with variable viral competence. In particular, this system can be used to understand how ecological attributes of host populations (e.g. density, diversity, immunity, heterogeneity) facilitate or impede emergence and how evolution proceeds as a virus adapts to a new host species (e.g. phenotypic changes, genetic changes, predictability, repeatability).

The *Caenorhabditis*-Orsay virus system joins a small set of empirical systems suitable for studying spillover and emergence. Prior studies using other systems have yielded useful insights into these processes. For example, bacteria-phage systems have been used to show that the probability of virus emergence is highest when host populations contain intermediate combinations of native and novel hosts [61], that pathogen variation in reservoir hosts drives emergence in novel hosts [62], and that mutations that allow phages to infect novel hosts also constrain further host range expansion [63]. Plant-virus systems have been used to document the effects of host species on the fitness distribution of viral mutations [64], to determine the importance of dose, selection, and viral replication for adaptation to resistant hosts [65], and to characterize how spillover can impact competition among host species [66,67]. *Drosophila-*virus systems have been used to show that viruses evolve in similar ways when passaged through closely related hosts [42] and to show that spillover dynamics can depend on temperature [68].

The *Caenorhabditis*-Orsay virus model can be uniquely useful for studying how ecology impacts spillover and emergence in animal systems since population characteristics like density, genetic variation, and immunity can be readily manipulated and virus transmission occurs without intervention by a researcher. *Caenorhabditis* hosts have complex animal physiology, immune systems, and behavior, meaning that this system can be useful for revealing the importance of variation in these traits. In this study, we identified multiple susceptible spillover hosts that have variation in transmission ability. In the future, these hosts can be used not only to probe how ecology impacts spillover and emergence, but also to better understand how and why spillover and emergence patterns may differ across hosts.

## ACKNOWLEDGEMENTS

We thank Marie-Anne Félix and Aurélien Richaud for sending *Caenorhabditis* strains and for advising on their propagation and on molecular species identification. We are also grateful to Marie-Anne Félix for her comments on an earlier version of this manuscript. We thank Anton Aluquin for help with viral extractions. We thank Beth Tuschhoff and Charles Geyer for helpful discussion about analysis and Andrew Wood for providing his expertise with Roar, the Penn State supercomputing cluster. We thank Lewis Stevens for technical guidance on working with phylogenetic data. We thank Amrita Bhattacharya, Heverton Dutra, Beth McGraw, and Andrew Read for lively discussion of spillover science and pattern interpretation. This work was partially supported by National Science Foundation grant DEB-1754692. The funders had no role in study design, data collection and analysis, decision to publish, or preparation of this article.

## Notes

### Competing Interest Statement

The authors have declared no competing interest.

