## Supplemental Information A for "Developing an empirical model for spillover and emergence: Orsay virus host range in *Caenorhabditis*"

Titration of virus filtrate

24-well plates were filled with 2 mL nematode growth medium (NGM) and seeded with 20 µL OP50 in LB broth [1]. OP50 lawns were allowed to grow at room temperature for approximately 24 hours. Eleven three-fold dilutions of the virus filtrate were made spanning concentrations of the original stock from 0.333 to 5.65 x 10^-6^. A total of 8 filtrate concentrations were used beginning with the 4^th^ of the dilution series (where the concentration of original filtrate was 0.01234), 20 µL diluted filtrate was added to the top of OP50 lawns in 6 wells.

*C. elegans* strain JU1580 and *C. briggsae* strain AF16 were bleach synchronized using standard methods [1]. 50 JU1580 or AF16 L1 were added, respectively, to each of 4 and 2 replicate wells for each virus filtrate dilution. Worms grew in wells for 3 days at 20°C. Worms were removed from the wells by pipetting 1 mL of water into and out of the wells. Worms suspended in water were transferred to microcentrifuge tubes. Worms were pelleted by centrifuging at 1000 x g for 1 minute, and then washed once by removing 900 µL of the supernatant and adding 900 µl fresh water. Tubes were again centrifuged to pellet worms. Supernatants were removed to 500 µL and the remaining worm – water mixture was moved to 2 mL round-bottomed snap cap tubes with approximately 100 µl 0.5 mm silica beads. These tubes were shaken in a TissueLyser II (Qiagen) for 2 minutes at 30 shakes per second. Tubes were then centrifuged at 17,000 x g for 5 minutes. The supernatant was transferred to a new microcentrifuge tube which was centrifuged again at 17,000 x g for 5 minutes. This supernatant was saved as the sample. Virus was quantified in samples by qPCR (see methods in text).

This assay assigns infection ability to a particular virus dilution based on whether there is more virus in wells containing JU1580 worms than AF16 worms (modified from [2]). Since none of the wells with AF16 worms had detectable virus in this assay, we classified JU1580 wells as infected or not based on the presence or absence of detectable virus (Figure 1). We used maximum likelihood to determine the number of infectious viral doses per 20 µL of the stock viral filtrate (analogous to the median tissue culture infectious dose, TCID50, [2]). In practice, this involved calculating the likelihood of observing the data for different values of TCID50 in R [3]. Code is available on GitHub. The values reported in the main text for the concentration of the stock viral filtrate are the maximum likelihood and the 95% confidence intervals determined as the range of TCID50 values within 1.92 log likelihood units of our maximum likelihood. We find that our stock viral filtrate is 8,562 (95% CI: 3,468 - 19,446) x TCID50 per 20 µL, which equates to 428.1 (95% CI: 173.4-972.3) x TCID50 per µL (Figure 2).

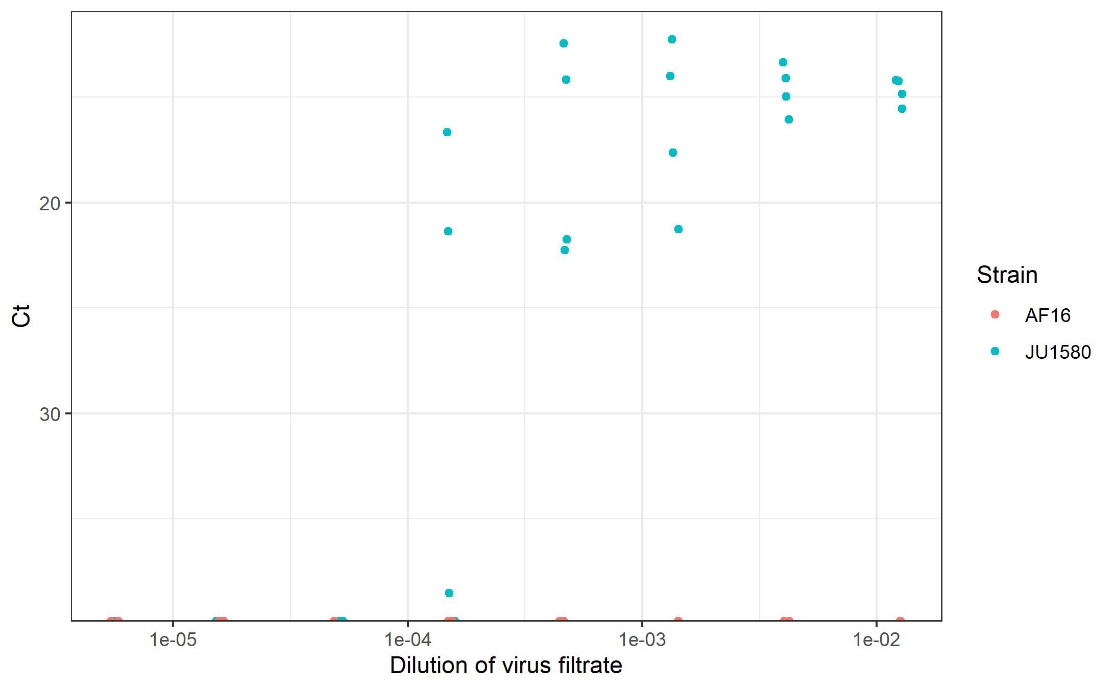

Figure 1. Quantification of infection across three-fold filtrate dilutions in *C. briggsae* AF16 (orange) and *C. elegans* JU1580 (teal). Points are slightly jittered on the x-axis to aid visualization.

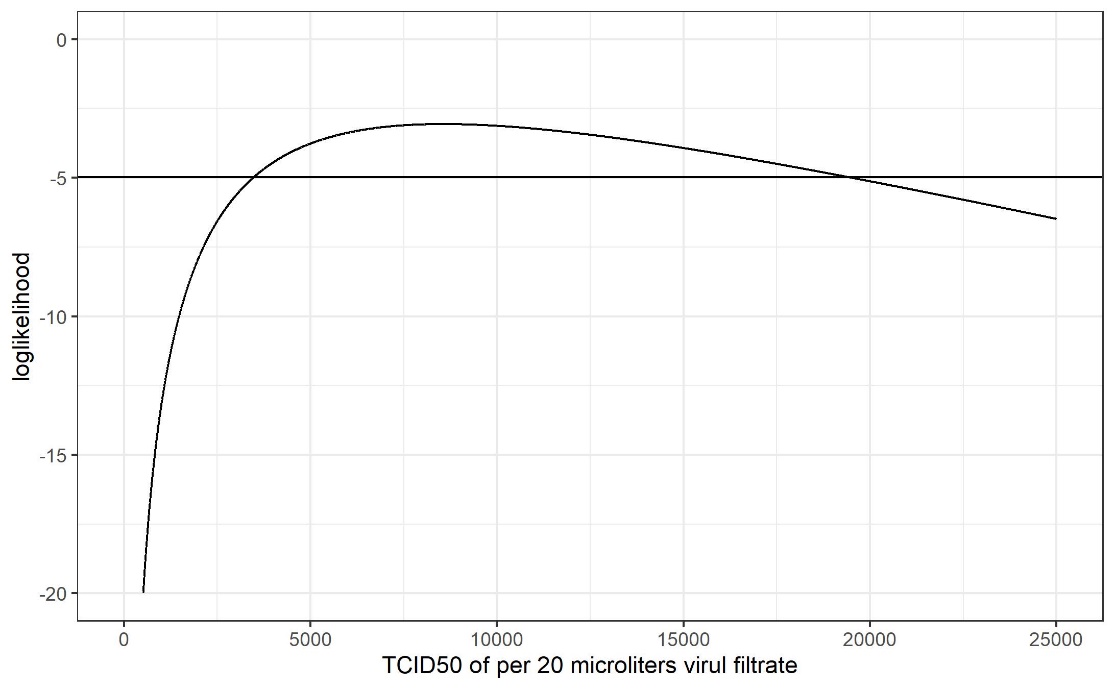

Figure 2. Log likelihood of TCID50 (per 20 µL) in the virus filtrate. The solid horizontal line is drawn at 1.92 log likelihood units below the maximum log likelihood. Values above this line thus depict all of the plausible values for TCID50.

3.	R Core Team. 2020 R: A Language and Environment for Statistical Computing.
